# Understanding the characteristic behavior of the wild-type and mutant protein structure of FLT3 protein by computational methods

**DOI:** 10.1101/2024.04.18.590047

**Authors:** Saleena Younus, Özge Tatli, Ahmad Nasimian, Julhash U. Kazi, Lars Rönnstrand

## Abstract

FLT3 emerges as a commonly mutated protein with significant prognostic implications in acute myeloid leukemia (AML). Point mutations or deletions in the tyrosine kinase domain (TKD) at the activation loop and internal tandem duplications (ITD) in the juxtamembrane (JM) region (and less commonly in the TKD) are the primary mutations that occur in the FLT3 protein. Besides, AML treatment with tyrosine kinase inhibitor drugs may result in the acquisition of TKD mutations in the FLT3-ITD structure. All these mutations will induce activation of the kinase activity of FLT3 protein leading to activation of downstream signaling pathways. Therefore, finding better therapeutics against each of these mutant FLT3 proteins is crucial in the treatment of AML. This study aims to comprehend the characteristic behavior of TKD mutants (C and F in Y842), ITD mutants, and the combination of ITD with TKD mutations (C and F in Y842) in the FLT3 protein through computational approaches, including Molecular Dynamic (MD) simulation, cluster analysis, and machine learning techniques. The MD simulation studies revealed the alterations in the optimized state, flexibility, and compactness nature between FLT3-WT and mutated FLT3 proteins and identified significant changes in the point mutants, ITD, and the combined ITD and TKD mutated FLT3 protein structures. Cluster analysis also confirmed that these mutations significantly impact the overall flexibility of the protein structures, especially in the point-mutated structures of FLT3-Y842C and FLT3-ITD-Y842F. These findings emphasize the diverse protein conformations of mutated structures of the FLT3 protein, contributing to the deregulation of FLT3 protein function, and identified these mutated proteins as promising therapeutic targets in the treatment of AML.

## Introduction

Acute myeloid leukemia (AML) is a type of leukemia, a disease that occurs in the blood and bone marrow, which is characterized as a heterogeneous hematological malignancy of hematopoietic stem cells [1, 2]. The prevalence of AML typically varies between the pediatric and adult cases based on age. In pediatric leukemia, AML accounts for approximately 15-20% of cases, while in adult leukemia, AML represents a higher proportion, exceeding 33% of leukemia cases [3]. Genetic mutation, use of chemotherapeutic drugs, and exposure to chemicals and radiation are the main underlying factors for the development of AML[4]. Also, multiple gene mutations have been associated with the development of AML. However, the mutation of FLT3, NPM1, CEBPA, RUNX1, ASXL3, and TP53 are considered as the more potent risk factors, in which FLT3, NPM1, and CEBPα have been thoroughly investigated in the treatment and disease progression of AML. The mutation rates of all these genes vary, and FLT3 is recognized as one of the most frequently mutated genes, correlating with a poor prognosis. The approximate mutation rate of FLT3 is around 30% in AML cases [5, 6]. In FLT3there are mainly two types of mutation found; one is the point mutation or deletion in the kinase domain (FLT3-TKD) of FLT3, which occurs in about 7-10% of AML. The other one is the internal tandem duplication mutation (ITD) in the JM region or TKD of FLT3, which is about 20-30% of all AML cases [7, 8]. The FLT3-ITD is an in-frame duplication of nucleotides that occurs in exons 14 and 15 of the juxtamembrane domain, which can vary in size between 3 up to ∼400bp nucleotides [9]. The ITD mutation of the FLT3 protein is the most aggressive and proliferative, with a high risk of relapse from the disease [10]. The FLT3-ITD mutations induce ligand-independent activation by disturbing the interaction between the juxtamembrane region and the kinase domain that normally keeps the kinase inactive [11],leads to autophosphorylation of FLT3 and activates multiple intracellular signalling pathways, including STAT5, MAPK, and AKT. Ultimately, these cascades leads to cell proliferation and anti-apoptosis [12, 13]. Additionally, TKD mutations may appear in the activation loop region of the FLT3-ITD mutated protein during treatment with tyrosine kinase inhibitors. The occurrence of those kinase mutations will likely result in the development of resistance to therapy in AML patients [14-16]. Developing more effective treatments targeting these mutants is thus of the utmost importance in the treatment of AML. To do this, we first need to understand the characteristic behaviour of wild-type (WT) and mutated FLT3 proteins.

In target-based treatment therapy, structure-based drug design (SBDD) techniques play a vital role in lead discovery and optimization. Currently, pharmaceutical companies and researchers are widely employing the SBDD method in their drug discovery and design process for identifying more specific and effective drugs for specific targets. The SBDD method consists of a collection of computational techniques, which includes structure-based virtual screening of the molecules (SBVV), molecular docking, and molecular dynamic (MD) simulation study of drug and drug-bounded protein complexes. These computational techniques reduce the time and cost of drug discovery and contribute to identification of more specific and effective drugs through high-throughput screening of small molecules [17, 18]. In SBDD, Molecular Dynamics (MD) simulation stands out as one of the most powerful computational techniques for elucidating the characteristic behaviour of the protein in biological systems. Using these techniques, we can simulate how the protein behaves during normal and diseased conditions. This information will give insightful knowledge of the biological behaviour of proteins at the molecular level of diseased pathways and further the drug discovery process. To enhance the therapeutic efficacy of drugs using SBDD, researchers are now incorporating new computational techniques, including machine learning, alongside MD simulation [19]. All these computational techniques will give further insightful information related to protein dynamics. Because understanding protein dynamics is crucial, as it directly correlates with conformational stability and flexibility, influencing the protein’s function and the cellular phenotype [20, 21].

In this study, our objective is to comprehend the characteristic behaviour of the FLT3 protein upon occurrence of various mutations, including (1) point mutations of the tyrosine residue Tyr842 (Y842) located in the kinase region in the activation loop to cysteine and phenylalanine residues (Y842C/F), (2) the internal tandem duplication (ITD) mutation (FLT3-ITD), and (3) a combination mutation of ITD in the juxtamembrane region and Y842C/F mutation in the kinase domain by computational techniques. The point mutation of C at the position 842 (TKD) in the FLT3-WT and the F mutation at the 842 position in the FLT3-ITD protein structure have already been reported [16, 22]. But the F mutation at residue 842 (TKD) in FLT3-WT and the C mutation at the 842 position in the FLT3-ITD mutated structure have not been reported. Through these investigations, we aim to gain a more comprehensive understanding of the characteristic behavior of this type of mutation in the FLT3 protein. This information will guide us to identify more effective therapeutics for AML treatment.

## 2. Materials and methods

For this study, we utilized the primary online resources were NCBI, NCBI-CDD, PDB, AlphaFold database, UniProt, VectorBuilder, and the SWISS-MODEL server [23-28]. Here the GROMACS software was employed for MD simulation studies of the protein structure [29]. The mutation of the structure was done with Pymol software, and visualization of the protein was done using Discovery Studio software [30, 31]. The generation of images for RMSD, RMSF, Rg H-Bond, and SASA were accomplished using Qtgrace (https://sourceforge.net/projects/qtgrace/) and Excel software. Cluster analysis of the structures, along with their visualization, was conducted through Rstudio software[32, 33]. The dynamic motion of both wild-type (WT) and mutated structures was visualized using Dynamut software [34].

### 2.1 Sequence comparison and protein modelling

For the study, we used the FLT3-ITD sequence data derived from AML patient and described earlier (the so-called W51 mutation[35]). This sequence has a 7 amino acids long duplication in the JM region. Based on this FLT3-ITD sequence, we performed multiple sequence comparisons of wild-type (WT) and FLT3-ITD protein sequences using tool from VectorBuilder website (https://en.vectorbuilder.com/tool/sequence-alignment.html). This tool shows 99.30% sequence identity of the FLT3-ITD sequence with the native FLT3 (WT) protein sequence and highlights the duplicated sequence regions of the FLT3-ITD protein. The details of the protein sequence alignment of FLT3-WT and FLT3-ITD are shown in the **Supplementary file S1**. From the protein sequence alignment of both WT and FLT3-ITD, we could identify that there is a duplication of seven amino acid residues **“REYEYDL”** occurring from 595-601 in the JM region of the FLT3-ITD sequence. Using this FLT3-ITD sequence, we modelled the protein structure of FLT3-ITD by homology modelling techniques using the SWISS–model server. Nowadays, several computational modelling software are available for the homology modelling of proteins. However, due to the easiness of handling software and considering the reliability of the modelled structures from the SWISS-model server, we used the SWISS-model server for modelling the FLT3-ITD structure.

### 2.2 Protein mutation

In the proposed work, we aimed to understand the characteristic effects in behavior by point mutation of Cysteine (C) or Phenylalanine (F) at position 842 in the activation loop region of the TKD of the FLT3 protein and identify the effect of the ITD (FLT3-ITD) mutation effect on the FLT3 protein. In this study, we are also trying to understand the effect of acquired mutations of either C or F in the Y842 position of the FLT3-ITD structure (Combination of ITD and TKD mutations). The acquired TKD mutations usually occur in the FLT3-ITD protein during the treatment of AML with TKIs [16, 22]. For this reason, in the FLT3-ITD structure, we made a point mutation in position 842 to C or F position. Here the effect of point mutation to C and F were analyzed in the WT and FLT3-ITD protein structure using Pymol software. In this study, we used six different structures of FLT3 protein, which included.,

1. FLT3-WT.
2. Point mutation structure of FLT3-WT with mutated residue in the TKD to Cysteine (Cys-C) in the activation loop region (FLT3-Y842C).
3. Point mutation structure of FLT3-WT with mutated residue in the TKD to Phenylalanine (Phe-F) in the activation loop region (FLT3-Y842F).
4. FLT3-ITD structure.
5. The structure of FLT3 proteins with a combination of FLT3 protein with ITD mutation and point mutation of C in position 842 (FLT3-ITD-Y842C).
6. The structure of FLT3 proteins with a combination mutation of FLT3 protein with ITD mutation and point mutation of F in position 842 (FLT3-ITD-Y842F).

Here we selected a modeled wild-type (WT) FLT3 protein structure from the Alphafold database (protein id; **P36888)**, which was generated using the Alphafold method. This choice was motivated by two factors: firstly, the structure closely mirrors the FLT3 protein sequence and does not exhibit any missing residues commonly found in X-ray crystallographic structures. Secondly, the modeled FLT3-ITD structure was selected based on this Alphafold-derived FLT3 protein structure.

### 2.3 Molecular dynamic (MD) simulation

Molecular Dynamic simulation is considered as one of the most powerful tools for understanding protein structure-related questions since it was first used in the study of rhodopsin protein in 1976 to understand its photoisomerization characteristics [36]. After that, the MD simulation study was considered as a vital thing for explaining the structure-function relationship of the proteins, especially the protein stability, function, protein-protein interaction, enzymatic reactions, protein-drug interaction, and membrane protein [37, 38]. Here GROMACS 2021 was employed for MD simulations of the FLT3 protein. For this study, we used six different structures of the FLT3 protein, including WT (FLT3-WT) with two different point mutated structures of WT (FLT3-Y842C and FLT3-Y842F), FLT3-ITD and combination mutation of ITD with two different TKD point mutated structures of FLT3-ITD structures (FLT3-ITD-Y842 and FLT3-ITD-Y842F). Before initiating the MD simulation run, we established the parameters for forcefield, solvation, energy minimization, and equilibration. The CHARMM27 force field was applied to the protein structure [39]. Subsequently, the entire structure was solvated within a triclinic box using the TIP3P water model [40]. To neutralize the entire system, a salt concentration of 0.15 M was maintained by adding the required number of Na^+^/Cl^-^ ions. The system’s energy was minimized with the steepest descent algorithm by 50,000 steps. A constant temperature of 300 K and 1.0 bar pressure were applied to the entire system based on the Berendsen algorithm and Parrinello-Rahman pressure-coupling to ensure equilibration at a constant temperature and pressure [41, 42]. In the MD simulation, Particle-Mesh-Ewald summation (PME) was utilized to calculate the electrostatic interactions of the system[43]. The NVT and NPT ensembles were applied for maintaining the equilibration and production of the entire system [44, 45]. The MD simulation was performed at 100 ns using the leap-frog integrator method [46]. In the MD simulation, 10,000 conformations were generated for the structure to analyze the trajectory. The trajectory of the MD simulation study was analyzed using built-in GROMACS tools, such as rms, rmsf, hbond, gyrate, and sasa[29].

### 2.4 Conformational stability of the wild-type and mutant protein by Cluster Analysis

To confirm the amino acid residue level flexibility, we utilized another computational technique, Cluster analysis, which is a widely used statistical technique to group similar objects or data points into clusters [32]. The main aim of cluster analysis is to partition a set of observations into subsets or clusters based on similarities between the data points within each group and dissimilarities between data points in different groups. Cluster analysis techniques now have been widely used in the structural biological studies of protein to understand the conformational changes of a protein by using the trajectories of MD simulation analysis of protein or protein complex. Cluster analysis uses different algorithms to divide the data set into clusters of similar properties. In the present work, we used the K-means algorithm to sort the objects from the data set into different groups. K-means indicate the sum of squared deviations (SSD) from the cluster’s centroid. By cluster analysis methods, we determined the optimal number of clusters in terms of RMSF using the elbow method. The “elbow” region of the objective function versus the number of cluster plots was sketched by R studio 4.2.2[33].

### 2.5 Evaluation of the stability of the wild and mutant of FLT3 protein by machine learning models

Apart from Gromacs software, we employed another software package, “Dynamut,” to confirm the dynamics of motion and stability of the wild-type and mutated structure of the FLT3 protein. Dynamut is a web-based server developed based on machine learning techniques (graph based signature) for assessing the impact of mutational changes of a protein structure and function relying upon static structures (wild type)[34]. Dynamut contains two tools for this purpose. One is the normal mode approach (NMA) to analyze and visualize protein dynamics motion by sampling the conformations of the protein structure, and another one is the “Prediction” tool to assess the impact of mutations on the protein dynamics and stability resulting from vibrational entropy changes. In the proposed work, we used the NMA approach to analyze the protein dynamic motion.

## 3. Results

### 3.1 Evaluation of the modelled protein

The 3D structure of the FLT3-ITD structure was obtained using the homology modeling technique of the SWISS-MODELLER Server. Using this server, we performed the template search of the FLT3-ITD sequence and then selected five template structures based on their sequence identity, GMQE, and coverage for building the FLT3-ITD structure. After running the build model, we selected the best model, which was based on the Swiss model template structure of FLT3 protein **P36888 (**https://alphafold.ebi.ac.uk/search/text/P36888) from the AlphaFold database. In the homology modelling server, the quality of the modelled structure was mainly evaluated by GMQE, sequence, identity, coverage, Molprobity, and Ramachandran plot (https://swissmodel.expasy.org/docs/help#model_evaluation). The GMQE (Global Model Quality Estimation) indicates the tertiary and quaternary structure level expected quality of the modelled protein, and its values usually range between 0 and 1 [47]. The highest value of the modelled structure indicates its higher expected quality. The GMQE score of the modelled protein structure was 0.75. The modelled structure shows 0.62 similarity and 100% sequence identity with the FLT3 protein developed by the AlphaFold method. The Ramachandran plot evaluated the energetically favoured regions for backbone dihedral angles against amino acid residues in the modelled protein structure. And the local and global levels of amino acid and protein level quality of the modelled structure were estimated by the MolProbity score. The low MolProbity score determines the best quality of the modelled structure. In the Swiss-Model server, both the Ramachandran and MolProbity score of the structure is estimated by MolProbity software (https://github.com/rlabduke/MolProbity)[48]. From the Ramachandran plot, we could identify that most of the amino acid residues that fall in the allowed region. The score of the favoured region was 90.08%, and the MolProbity score of the modelled structure was 1.50. Based on these values, we can say that the modelled protein structure is stereo-chemically more stable. The details of the selected template structure(A), best model structure, and their GMQE, sequence identity(B), Ramachandran plot(C), and 3D structure of the best model protein and their ITD mutated structure(D) are shown in **Figures 1**.

**Figure 1:**
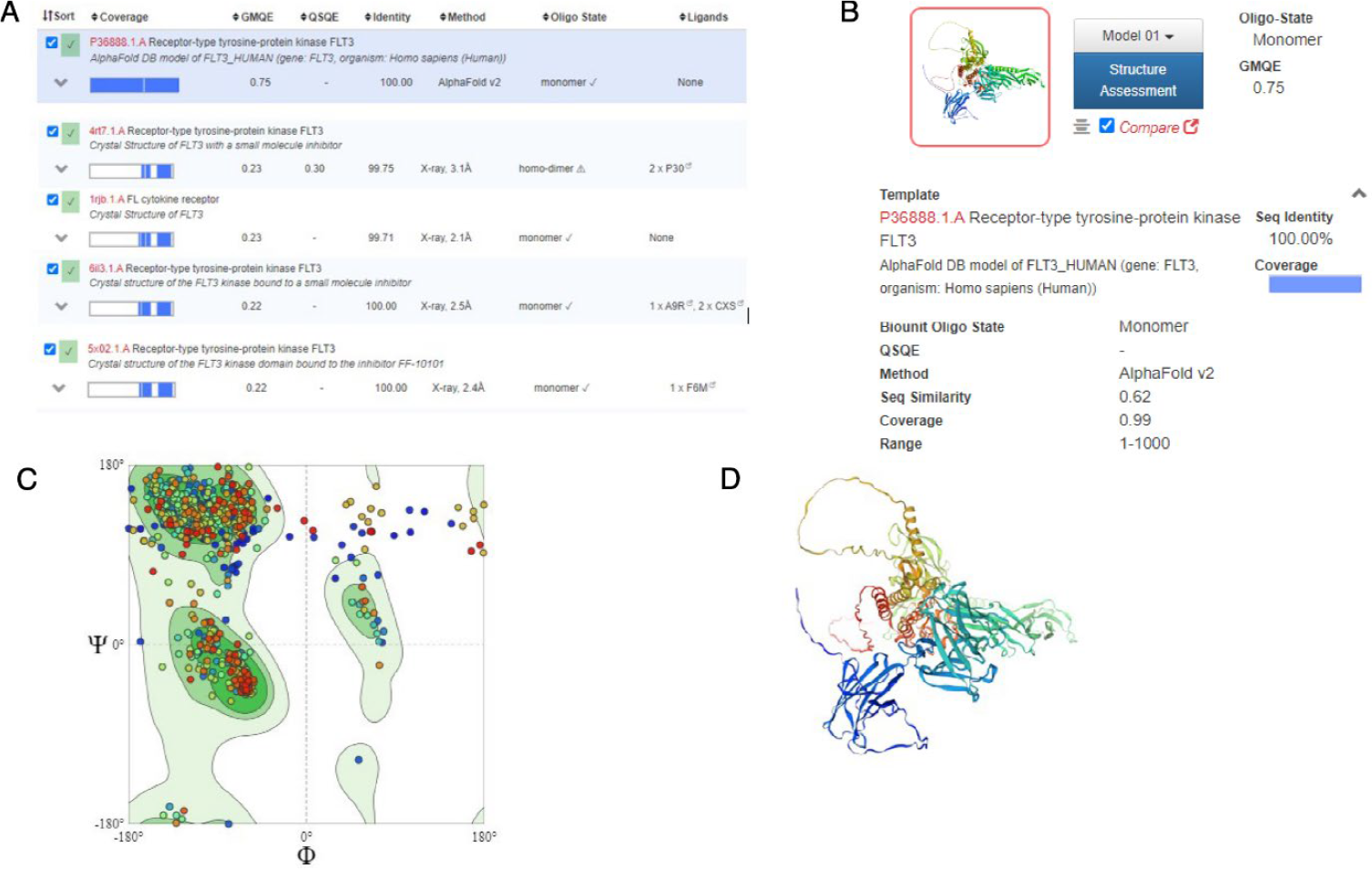
**(A)** Details of the five selected template structures of FLT3 protein used for the homology modeling of the FLT3-ITD protein structure. **(B)** Details of the final modeled structure selected from the built structure by homology modeling. **(C)** Ramachandran plot of the selected structure. **(D)** 3D structure of the modelled protein structure FLT3-ITD.

### The structural details of FLT3-WT and FLT3-ITD protein structure

The FLT3-WT protein structure is composed of a total of 993 amino acid residues with a molecular weight of about 150-160 kDa. The protein structure encompasses an extracellular N-terminal region, a transmembrane region (TM), a juxtamembrane region (JM), and an intracellular C-terminal region. The extracellular N-terminal region includes five immunoglobulin-like domains (D1 (27-161), D2 (162-244), D3 (245-346), D4(347-434), and D5(435-541)), encompassing amino acid residues from 27 to 541. Among these, three domains are involved in ligand binding, and two are involved in protein dimerization. The extracellular region consists entirely of beta-strand structures. The TM domain adopts a helical structure composed of amino acid residues 544-563, anchoring the protein in the cytoplasmic region. The 564-571 region is considered a linker [11]. The juxtamembrane domain (JM) (comprising amino acid residues 572-603) controls the FLT3 protein activation by steric inhibition and maintains the protein in an inactive form. The JM region consists of three sub-motifs: the JM binding motif (JM-B, 572-578), the JM switch motif (JM-S, 579-592), and the JM zipper motif or linker motif (JM-Z, 593-603). The intracellular kinase domain (amino acid residues 604-958) is highly conserved. Within the kinase region of the FLT3 protein, there are two substructures, an N-lobe and a C-lobe, containing two sub-kinase domains (TKD1 and TKD2) which are interconnected by another kinase substructure known as the activation loop (A-Loop, amino acid residues 829-858). In the modelled FLT3-ITD protein structure includes 1000 amino acids with continuous numbering, the duplicated region appears from positions 595 to 601, while the original position of the seven amino acid residues is numbered as positions 602 to 608 in the JM. Our study focused on the Y842 amino acid residue in both FLT-WT and FLT3-ITD. For maintaining consistent numbering in the FLT3-ITD protein structure as in FLT3-WT, we revised the entire numbering of the FLT3-ITD structure and adjusted the numbers of both the duplicated region and the original region of the JM-Z motif with the same numbering **(“REYEYDL**” 595-601). The detailed entire 3D structure view of FLT3-WT and FLT3-ITD (**Figure 2A and Figure 2B**) and its super imposed views (**Figure 2C, Figure 2D**) are shown below.

**Figure 2:**
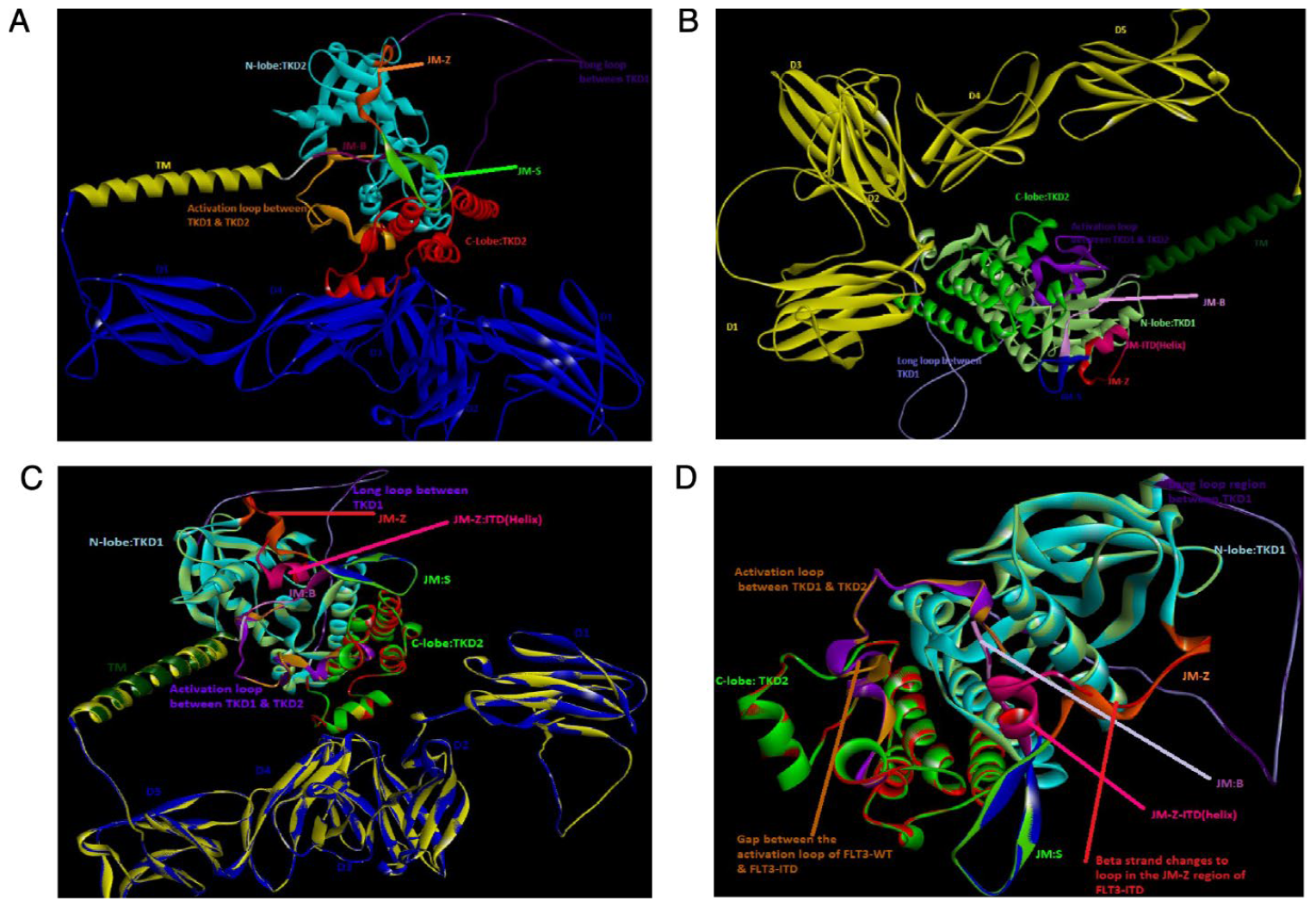
Full 3D view of the protein structures. (A) FLT3-WT. (B) FLT3-ITD. (C) Full 3D view of superimposed view the protein structures FLT3-WT and FLT3-ITD. (D) Superimposed view of JM and kinase domain of FLT3-WT and FLT3-ITD.

For the computational study, we only considered the JM and kinase parts of FLT3-WT and FLT3-ITD (572-958). In the FLT3-ITD structure, duplication occurred in the JM-Z region. The JM-Z region comprises two small beta strands, which are interconnected by a small loop region (11 amino acid residues from 593 to 603). When examining the superimposed structure in **Figure 2d**, it’s evident that the duplication of the pattern “REYEYDL” forms two small helical patterns in the JM-Z region of FLT3-ITD and also the first strand completely converted into a long loop. These conformational changes may be the main reason for the loss of the steric inhibition nature of the JM region in FLT3-ITD, leading to the protein being in an active form. In FLT3-WT, the activation loop has a small folded nature in the mid-portion. But in the superimposed structure, this folded nature has been altered by ITD duplication. These changes will make the activation loop more flexible and increase the autophosphorylation rate of the FLT3-ITD protein.

### 3.2 MD simulation

Protein functionality is directly correlated with its conformational stability and flexibility [34, 36, 37]. This type of information will be helpful for understanding the nature of different mutations and their impact on the diseased pathway of AML. Based on this information, we can find more effective therapeutics against target proteins to control deregulation of pathophysiological pathways. In this study, our primary objective was to elucidate the characteristic behaviours of FLT3 protein with distinct mutations, including point mutations with C and F amino acid residues at position 842 in the activation loop of the TKD (Y842C/F), ITD mutation (FLT3-ITD), and the combination mutation of ITD with C and F point mutations at position 842 in the FLT3 protein structure (FLT3-ITD-Y842C/F). Here we mainly analyzed the trajectories including RMSD, RMSF, Radius of Gyration (Rg), H-Bond and SASA by using inbuilt tools of Gromacs. To explore the variations of FLT3-WT and mutated structure, we show three types of plots of RMSD, RMSF, Rg, SASA, and H-Bond, which included the plots of.,

- FLT3-WT versus FLT3-Y842C and FLT3-Y842F.
- FLT3-WT versus FLT3-ITD, FLT3-ITD-Y842C and FLT3-ITD-Y842F (combination mutation structure).
- FLT3-WT versus FLT3-Y842C, FLT3-Y842F, FLT3-ITD, FLT3-ITD-Y842C and FLT3-ITD-Y842F.

#### a. Root Mean Square Deviation (RMSD)

RMSD values are used for analysing the structural deviation of a protein. Usually, less RMSD values are considered for good structures to evaluate their structural stability [49]. Here, we performed the RMSD analysis for the six different structures of FLT3 protein, which are shown in three different plots., **A, B, C** in **Figure 3**. Plot **A** contained the comparison plots of FLT3-WT with FLT3-Y842C and FLT3-Y842F point mutation structures and **B** contains the RMSD plots of FLT3-WT with FLT3-ITD and combination mutation structures of FLT3-ITD-Y842C and FLT3-ITD-Y842F (point mutation of C and F in position 842 of the ITD structure). Plot **C** contained FLT3-WT and all the mutated structures of FLT3 protein. In **Figure 3A**, RMSD plot, we can see that in 12.5ns, the movement of FLT3-Y842C shows higher within the range 0.4nm-1.25nm, and in 50ns, it showed the deviation range almost near to 1.2nm. After 60 ns, the FLT3-Y842C mutated structure shows almost a constant deviation within the range of 0.78nm. During the range of 12.5 ns – 50 ns, FLT3-Y842F underwent to a slightly higher deviation within the range of 0.75 nm-1nm. Following this, the deviation becomes constant near to 0.6nm, and this deviation continues until 100ns. During 70-75ns, the deviation of FLT3-WT as within the range of .75nm and 1nm. Besides, the deviation of FLT3-WT was comparatively less than that of FLT3-Y842C and FLT3-Y842F. The FLT3-WT with FLT3-ITD, FLT3-ITD-Y842C and FLT3-ITD-Y842F plots of RMSD in **Figure 3B**., **FLT**3-ITD-Y842F showed a slightly higher deviation than all the other structures. Apart from this deviation, all the structures maintained a deviation within the range of 0.5nm and 1nm. However, when compared all the mutated structures RMSD plots with FLT3-WT in **Figure 3C**, we can see that the deviation range of FLT3-WT is different than its mutated structures. Here, we can clearly understand that the rate of deviations of FLT3-Y842C, FLT3-Y842F and FLT3-ITD-Y842F mutated structures shows higher when compared to all other structures. The average RMSD values of FLT3-WT, FLT3-Y842C, FLT3-Y842F, FLT3-ITD, FLT3-ITD-Y842C and FLT3-ITD-Y842F were 0.71 nm, 0.80 nm, 0.79 nm, 0.73 nm, 0.72 nm and 0.75 nm respectively.

**Figure 3:**
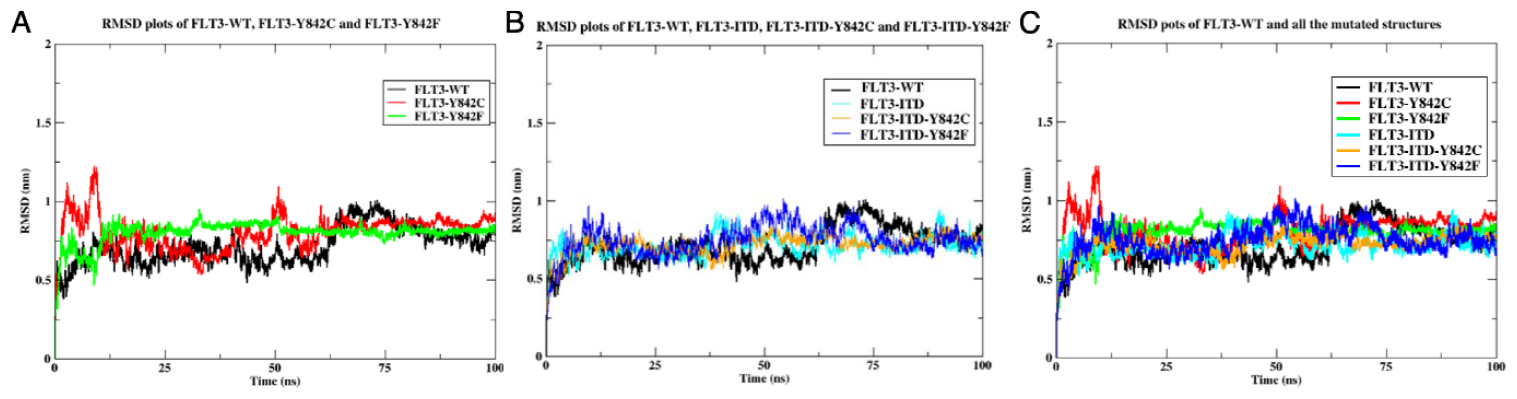
RMSD plots of WT and mutated structures of FLT3 protein. (**A)** RMSD plots FLT-WT with FLT3-Y842C and FLT3-Y842F (**B)** is the plots of FLT3-WT with FLT3-ITD, FLT3-ITD-Y842C and Y842F respectively. **(C)** Combined RMSD plots of WT with FLT3-Y842C, FLT3-Y842F, FLT3-ITD, FLT3-FLT3-ITD-Y842C and FLT3-ITD-Y842F.

#### b. Root mean square fluctuation (RMSF)

RMSF values are used to understand a protein’s overall amino acid residue level fluctuations at a given time. This information will be useful for understanding the folder nature of the protein [50, 51]. In a protein structure, the loop and terminal regions have more chances to fluctuate. When we look at the RMSF1 plots of FLT3-WT (**Figure 4A**) with point mutated structures of FLT3-Y842C and FLT3-Y842F, we can see that the fluctuation rate of loop (in which is located between 730-769) and terminal region of FLT3-Y842C are comparatively higher than FLT3-WT and the fluctuation rate of the loop and terminal regions of FLT3-Y842C went near to 1.70nm and 0.75nm respectively. From the plots, we could also identify that when F mutation occurs in the FLT3-WT, the fluctuation rate of the entire amino acid residues has decreased (FLT3-Y842F). This indicates that the native conformation of the FLT3-WT protein has been changed by the C and F mutation, and the FLT3-Y842C underwent to a more flexible conformation compared to FLT3-WT. In contrast, FLT3-Y842F structure acquired a rigid form than FLT3-WT. When we look at the RMSF (**Figure 4B**) plots of FLT3-WT with mutated structures of FLT3-ITD, FLT3-ITD-Y842C, and FLT3-ITD-Y842F, we can identify that, here the entire fluctuations of the amino acid residues of the mutated structures have been changed included loop and terminal region than FLT3-WT. The fluctuation rate of the loop region of FLT3-ITD-Y842F shows comparatively higher fluctuations and going up to 1.4nm. Here, the fluctuation rate of FLT3-ITD-Y842F was higher than FLT3-WT, and other mutated structure fluctuations were below than FLT3-WT. This means the FLT3-ITD-Y842F structure is more flexible than FLT3-WT, and FLT3-ITD and FLT3-ITD-Y842C are more rigid than FLT3-WT. When we look at the overall fluctuation rate of all the six structures in **Figure 4C**), we can identify that the FLT3-Y842C and FLT3-ITD-Y842F mutated structure went to more flexible than FLT3-WT and other mutated structure goes to more rigid than FLT3-WT. The average RMSF values of FLT3-WT, FLT3-Y842C, FLT3-Y842F, FLT3-ITD, FLT3-ITD-Y842C and FLT3-ITD-Y842F were 0.25nm, 0.33nm, 0.18nm, 0.23nm, 0.20nm and 0.29nm respectively.

**Figure 4:**
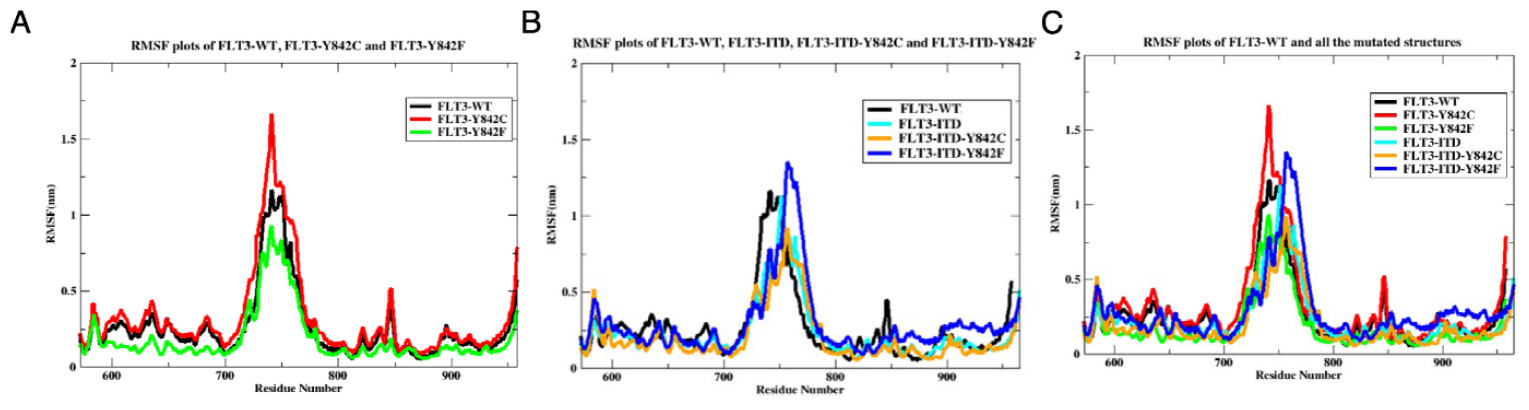
RMSF plots of WT and mutated structures. **(A)** RMSF plots of FLT3-WT, FLT3-Y842C and FLT3-Y842 structures. **(B)** RMSF plots of WT and combination mutation of ITD and TKD (C and F point mutation) structures. FLT3-WT, FLT3-ITD, FLT3-FLT3-ITD-Y842C and FLT3-ITD-Y842F. **(C)** Combined RMSF plots of WT with FLT3-Y842C, FLT3-Y842F, FLT3-ITD, FLT3-FLT3-ITD-Y842C and FLT3-ITD-Y842F.

#### c. Radius of gyration (Rg)

Rg is used for assessing the total compactness nature of a protein structure and estimating the compactness in the X, Y, and Z axes as a function of time when it moves (https://manual.gromacs.org/2019/onlinehelp/gmx-gyrate.html) [52]. From the assessment, we can determine how the dimension and shape of the structure vary when it undergoes different mutations. Here, we also considered the low Rg value for assessing the good compactness of the protein. As depicted from **Figure 5A**, the FLT3-Y842C shows slightly higher deviations between 0ns and 20ns and in 50ns within the range 2.6nm and 2.5nm, respectively, when compared with FLT3-WT. Except for these two high variations, FLT3-Y842C shows a high compactness nature until 100 within the range of 2.25nm. Also, the FLT3-Y842F structure shows a good compactness nature when it comes to a point mutation in the FLT3-WT with F mutation in position 842, and its Rg values became 2.25nm until 100ns after 20ns. Upon examining the atomic movements along the X, Y and Z axes of FLT3-WT, FLT3-Y842C, and FLT3-Y842F, it becomes the evident that, the atom movements of X and Z of FLT3-Y842C structure are slightly higher in the time 20ns and also between 40-60ns within the range 2.25nm-2.5nm. Apart from this deviation, all the structures’ X, Y, and Z direction atom movements are between 1.5nm-2.25nm. From the different compactness nature of FLT3-Y842C and FLT3-Y842F, we can say that the FLT3-Y842C structure is more flexible and FLT3-Y842F is more rigid than its native structure from FLT3-WT. When we compared the Rg2 plots (**Figure 5B**) of FLT3-WT with FLT3-ITD, FLT3-ITD-Y842C, and FLT3-ITD-Y842F, we could identify that FLT3-ITD and FLT3-ITD-Y842F structure have some high deviation in their structure compactness within the range 2.25nm-2.5nm in between 10ns and 70ns-100ns. In the case of FLT3-ITD-Y842C, we can see that this structure attained a good compactness nature within the range of 2.20nm-2.25nm after 10ns. Here, we can also see that atoms’ movements in all the structures’ X, Y, and Z directions are within the range of 1.6nm-2.25nm. From these plots, we could identify the flexibility and rigidity of the mutated protein. When we compared the overall compactness nature of the mutated structures of FLT3 with FLT3-WT, Rg3(**Figure 5C**), we could identify that FLT3-Y842C, FLT3-ITD, and FLT3-ITD-Y842F, which shows less compactness (more flexible) in the protein structure and other mutated structure show high compactness nature(rigid) than WT. The average Rg values of FLT3-WT, FLT3-Y842C, FLT3-Y842F, FLT3-ITD, FLT3-ITD-Y842C, and FLT3-ITD-Y842F were 2.32nm, 2.33nm, 2.24nm, 2.36nm, 2.26nm and 2.33nm respectively.

**Figure 5:**
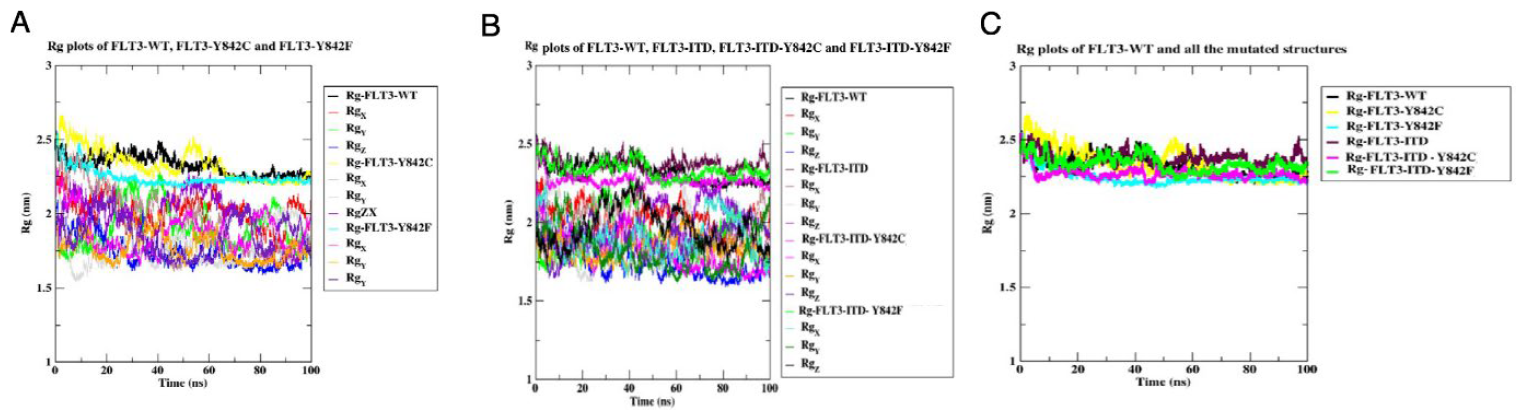
Rg plots of WT and mutated structures of the FLT3 protein. Here the upper region plots indicate the total compactness nature and lower plots indicate the x, y and z direction movements of atoms in the protein with respect to time. **(A)** The total compactness nature of FLT3-WT, FLT3-Y842C and FLT3-Y842 structures indicated by the black, yellow and cyan indicate. The lower plots indicate the X, Y and Z axis movements of atoms in each protein structure with respect to time. **(B)** Total Rg (black, maroon, magenta and green) and X, Y and Z direction movement of atoms within the protein structures, FLT3-WT, FLT3-ITD, FLT3-FLT3-ITD-Y842C and FLT3-ITD-Y842F. **(C)** Combined Rg plots (total compactness) of WT with FLT3-Y842C, FLT3-Y842F, FLT3-ITD, FLT3-FLT3-ITD-Y842C and FLT3-ITD-Y842F respectively.

#### d. Solvent Accessible Surface Area (SASA)

SASA of a protein is used for estimating the surface area of protein that is directly accessible to the surrounding solvent molecule in water. Based on SASA plots, we can predict a protein’s dynamic stability and behaviour in solution [53-55]. The lower SASA value indicates that the greater folding stability of the protein and also the hydrophobic residues of a protein are more exposed. The comparison of SASA plots of FLT3-WT with FLT3-Y842C and FLT3-Y842F reveals that, in between 40ns-60ns SASA of FLT3-Y842C was slightly vary than FLT3-WT structure and its value went to between 230nm-240nm **(Figure 6A**). Thereafter, it exhibits a better-folded state than FLT3-WT. In FLT3-Y842F, which shows slight variation between 0ns-20ns, and its SASA value ranges between 220nm-240nm; after that, its SASA value decreased in between 200nm-220nm, and this variation remained a constant at 100ns. In both variations, we can identify the flexibility and rigidity of the FLT3-Y842C and FLT3-Y842F mutated structures in water. In The SASA plots of FLT3-WT with mutated structures FLT3-ITD, FLT3-ITD-Y842C, and FLT3-ITD-Y842F (in **Figure 6B**) reveals that, the mutated structures goes to instability in their folded nature. Initially, the FLT3-ITD-Y842F surpassed 250nm within 10ns, but thereafter, it dropped to a lower value. However, in the case of FLT3-ITD, FLT3-ITD-Y842C, and FLT3-ITD-Y842F structures, we can see that its SASA value is increasing than FLT3-WT from 35ns and SASA value ranges in between 220nm-240nm and also this unfolded nature kept at 100ns. Based on the comparison SASA plots of all the six structures (**Figure 6C**), it can be discerned that the mutated structure FLT3-Y842F exhibits higher folded nature than WT, whereas the remaining mutated structures demonstrate less folded nature than WT in solution. The average SASA values of FLT3-WT, FLT3-Y842C, FLT3-Y842F, FLT3-ITD, FLT3-ITD-Y842C, and FLT3-ITD-Y842F were 218.62nm, 216.48nm, 211.69nm, 227.21nm, 223.59nm and 225.58nm respectively.

**Figure 6:**
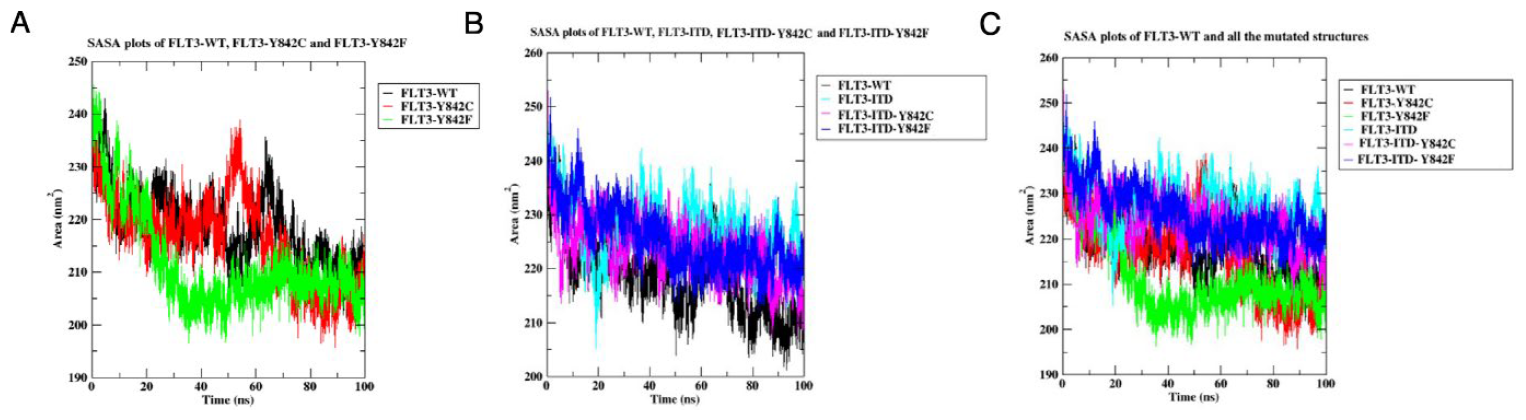
SASA plots of WT and mutated structures of FLT3 protein. **(A)** SASA plots of WT and mutated structures of FLT3-WT, FLT3-Y842C and FLT3-Y842F. **(B)** Wild and combination mutation structures of FLT3-WT, FLT3-ITD, FLT3-ITD-Y842C and FLT3-ITD-Y842F of FLT3 protein. **(C)** Comparison SASA plots of FLT3-WT with FLT3-Y842C, FLT3-Y842F, FLT3-ITD, FLT3-FLT3-ITD-Y842C and FLT3-ITD-Y842F respectively.

#### e. Intra Hydrogen Bond (H-Bond) interaction

Intra-molecular hydrogen bond interaction between amino acid residues in a protein plays an important role in maintaining its conformational stability, which was first proven in the 1980s by the Fersht group [56, 57]. After their discovery, several studies were conducted to understand the importance of intra-molecular hydrogen bond interaction between amino acid residues in a protein and have reached similar conclusions. Here, we try to find the intra-molecular hydrogen bonding stability of FLT3-WT with mutated structures of FLT3 protein by MD simulation. A higher H-Bond interaction reveals that the protein conformational stability is very good. Here, from the graph of H-Bond interaction plots of FLT3-WT with point mutated structure of FLT3-Y842C and FLT3-Y842F in **Figure 7A**, the FLT3-Y842C shows a higher H-bond interactions between 15ns-17ns within the 250-300 range and another in the 85-95ns within the 300 range and also in FLT3-Y842F structure, the number of intra-molecular H-Bond interaction increased to 300range within 40-60ns. Here, we could identify the bonding instability of both the mutated structures. In **Figure 7B** H-Bond interaction plots reveals the intra H-Bond interaction of FLT3-WT with ITD and the combination mutation of ITD and TKD of FLT3 protein. Here the intra H-bond interactions was slightly greater in point mutated structures of FLT3-ITD protein structure, when comparing the FLT3-WT with ITD and the combined mutation of ITD and TKD of FLT3 protein, its ranges exceeded 300 in between 40ns-100ns. Upon evaluating the overall comparison study of FLT3-WT with point mutation, ITD and combination mutation plots of **Figure 7C**, it become evident that the combination mutation structure of FLT3-ITD-Y842C and FLT3-ITD-Y842F exhibits a high number of intra-molecular H-bond interactions when compared to all other structures of FLT3. From the entire comparison study of H-Bond interaction, we can conclude that protein conformation has been changed, when mutation introduced in the FLT3 protein. The average number of intra-molecular H-Hond interactions in FLT3-WT, FLT3-Y842C, FLT3-Y842F, FLT3-ITD, FLT3-ITD-Y842C, and FLT3-ITD-Y842F were 274, 270,276,271,280 and 278 respectively.

**Figure 7:**
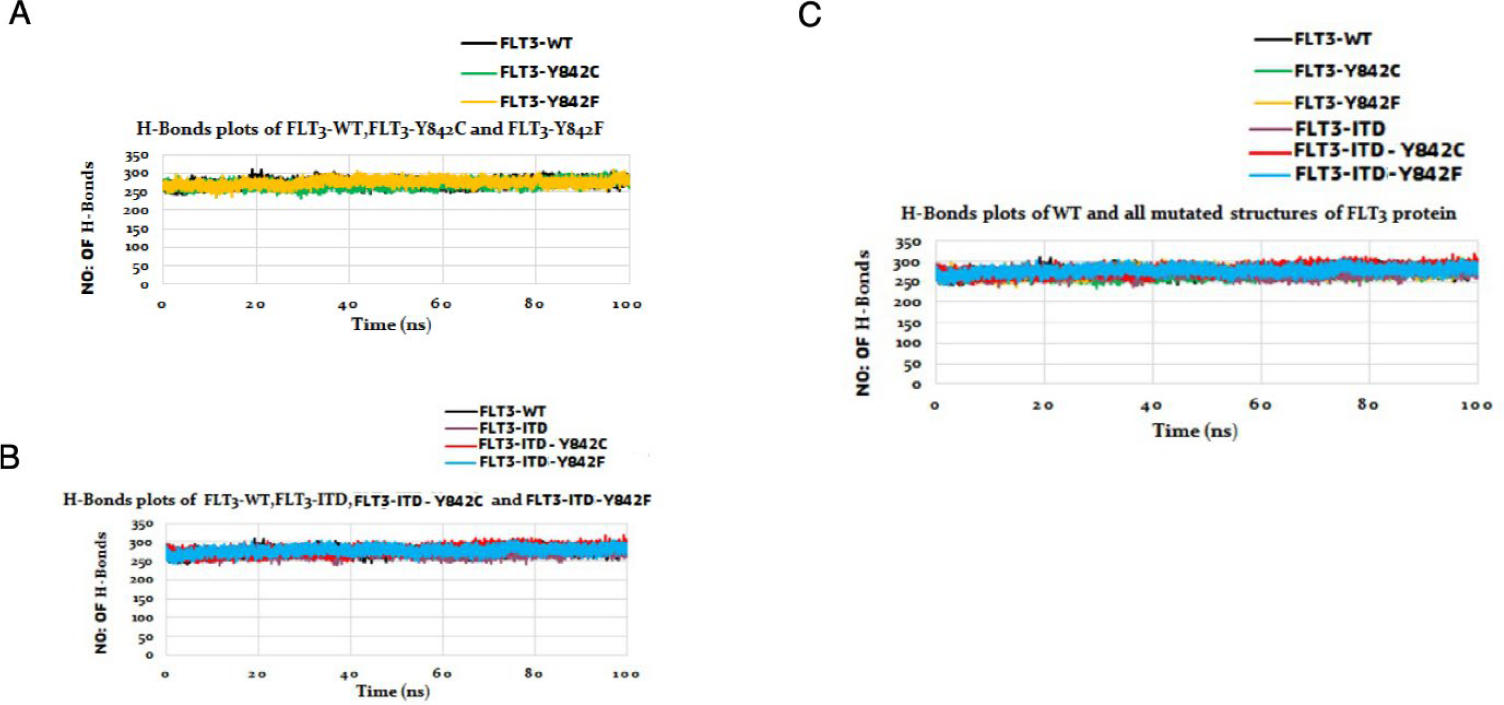
Intra-H-Bond interaction plots of WT and mutated structures of FLT3 ptotein. **(A)** H-Bond interaction graph of FLT3-WT, FLT3-Y842C and FLT3-Y842F of FLT3 protein. **(B)** FlT3-WT and combination mutation structure FLT3-ITD-Y842C and FLT3-ITD-Y842F structures of FLT3 protein. **(C)** The intra-H-Bond plots of WT with FLT3-Y842C, FLT3-Y842F, FLT3-ITD, FLT3-FLT3-ITD-Y842C and FLT3-ITD-Y842F, respectively.

### 3.3 Cluster Analysis

To understand the conformational states of the WT and mutated structures, we used the protein backbone trajectories from the RMSF data. Based on the cluster analysis study, we can group the similar nature of the amino acid residue of the protein. For grouping the data, we used the K-means clustering algorithm, and then optimal number of clusters identified by elbow method from the RMSF data of FLT3-WT, FLT3-Y842C, FLT3-Y842F, FLT3-ITD, FLT3-ITD-Y842C, and FLT3-ITD-Y842F. In **Figures 8(A-F)** represent the elbow graph of WT and mutated structures of FLT3 proteins. In the elbow graph, the X-axis represents the number of clusters, and the Y-axis represents the total within the sum of squares. Plots in **Figure 8 (B, D, F, H, J, and L)** represent the pictorial representations of cluster plots of WT and mutated structures of FLT3 protein structure based on the elbow graph. Here, the X-axis represents the normalized amino acid residue ranges, and the Y-axis represents the deviations (RMSF) of the amino acid residues. When we look at the elbow graph of WT and mutated structures of FLT3, we could identify that the maximum three normalized clusters from FLT3-WT, FLT3-Y842C, FLT3-ITD, FLT3-ITD-Y842C (**Figure 8 (A, C, G, and I)**) and four clusters from FLT3-Y842F and FLT3-ITD-Y849F (**Figure 8 (E, K**) can generate from the RMSF data. Based on this, we generated the appropriate cluster plots (**Figure 8 (B, D, F, H, J, and L))** by RStudio. From the cluster plots, we can see that, in all the protein structures, the majority of the amino acid residues fall under zero, and also, the lowest part of the clusters are more crowded than in all the cluster plots. From the secondary structure analysis of all the structures by Pymol software, we could identify that in all the structures, helix and sheets fell in the lowest part of the clusters, and the loop regions went above the upper part. From the detailed analysis of number of amino acid residues in the upper and lower part of the clusters from each structure details, we could identify that total number of amino acid residues fell below zeros in the FLT3-WT, FLT3-Y842C, FLT3-Y842F, FLT3-ITD, FLT3-ITD-Y842C and FLT3-ITD-Y842F were 223, 176,327,270,286 and 202, respectively.

**Figure 8:**
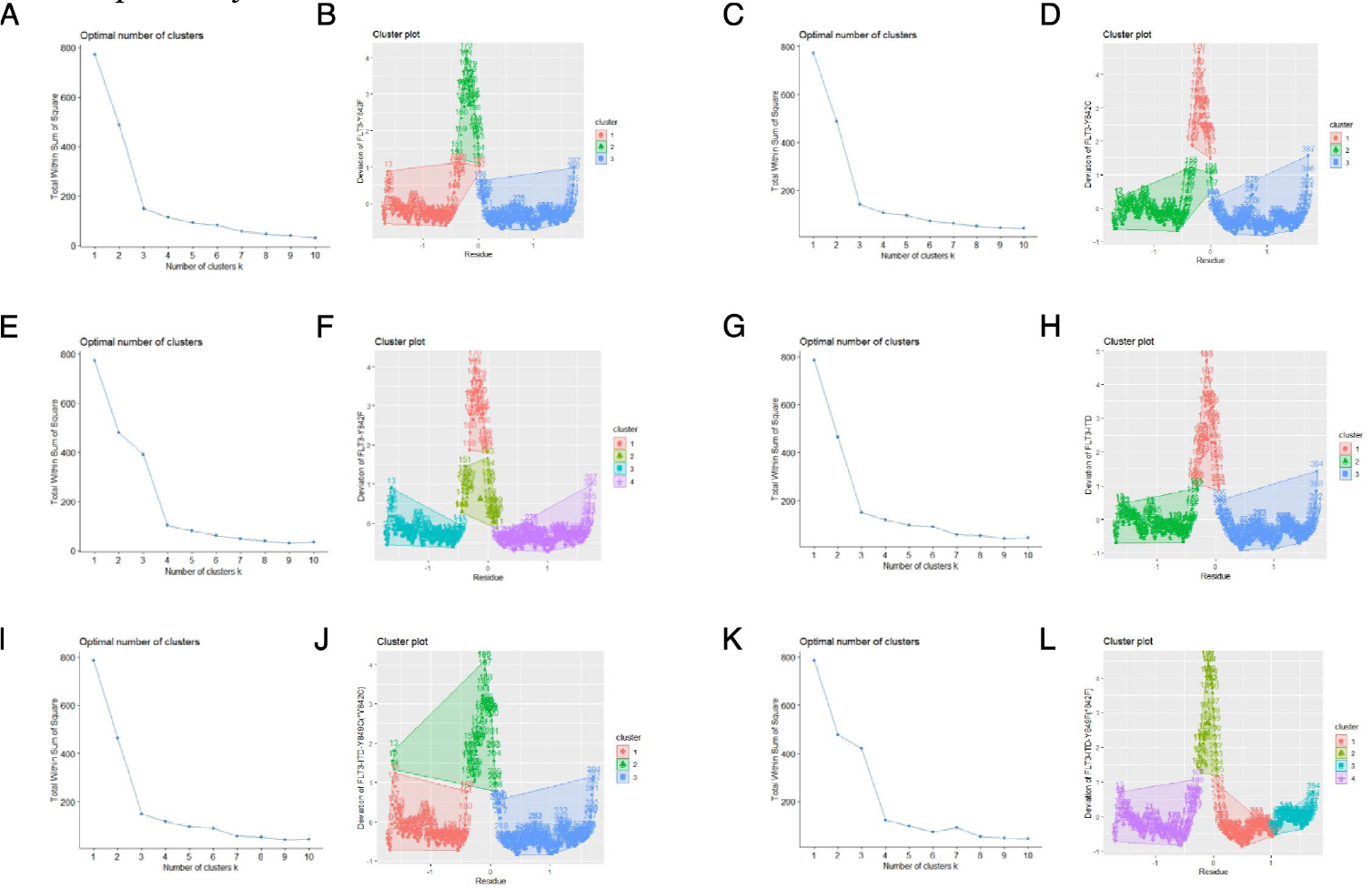
Elbow and Cluster plots of WT and mutated structures of FLT3-WT, FLT3-Y842C, FLT3-Y842F, FLT3-ITD, FLT3-ITD-Y842C and FLT3-ITD-Y842F of FLT3 protein based on RMSF data.

### 3.4 To understand the protein dynamic movements of the WT and mutant type by Machine learning tools

To understand the dynamic behaviours (motion of the protein structure) of the WT and FLT3-ITD structures, we performed NMA analysis for structures. The NMA uses a consensus method DUET that combines the statistical potential function for energy calculations (SDM), structural pattern mining approach (mCSM-stability), and ENCoM, including the nature of the protein from the structural information of a natively folded protein to calculate the motion of the protein structure. Using DUET and ENCoM, NMA analyses the approximate dynamics of a system around a conformation through harmonic motion. By using this harmonic motion, it will generate possible movement. From this movement, we can gain valuable insight into protein motions. **Figure 9** represents the harmonic movement (A, C, E, G, I, and K) and their steady-state (B, D, F, H, J, and L) images of the movement direction of WT and mutant type FLT3 protein structure by NMA analysis. From the molecular motion of FLT3-WT, FLT3-Y842C, FLT3-Y842F, FLT3-ITD, FLT3-ITD-Y842C and FLT3-ITD-Y842F, we could identify that the trajectory motion of the FLT3-Y842F, FLT3-ITD, FLT3-ITD-Y842C and FLT3-ITD-Y842F are in the opposite direction than the movement of FLT3-WT and FLT3-Y842C. In WT FLT3 protein, the movement starts from the downward direction and then goes to the upward direction. In the case of the FLT3-Y842C mutated structure, the movement direction is the same as FLT3-WT. However, in the case of FLT3-Y842F, FLT3-ITD, FLT3-ITD-Y842C, and FLT3-ITD-Y842F mutated structure, the movements start from the upward direction after its movements go in a downward direction. Trajectory representation for the first non-trivial mode of the molecule motion based on the Normal Mode Analysis are shown in the **Figure 9**.

**Figure 9:**
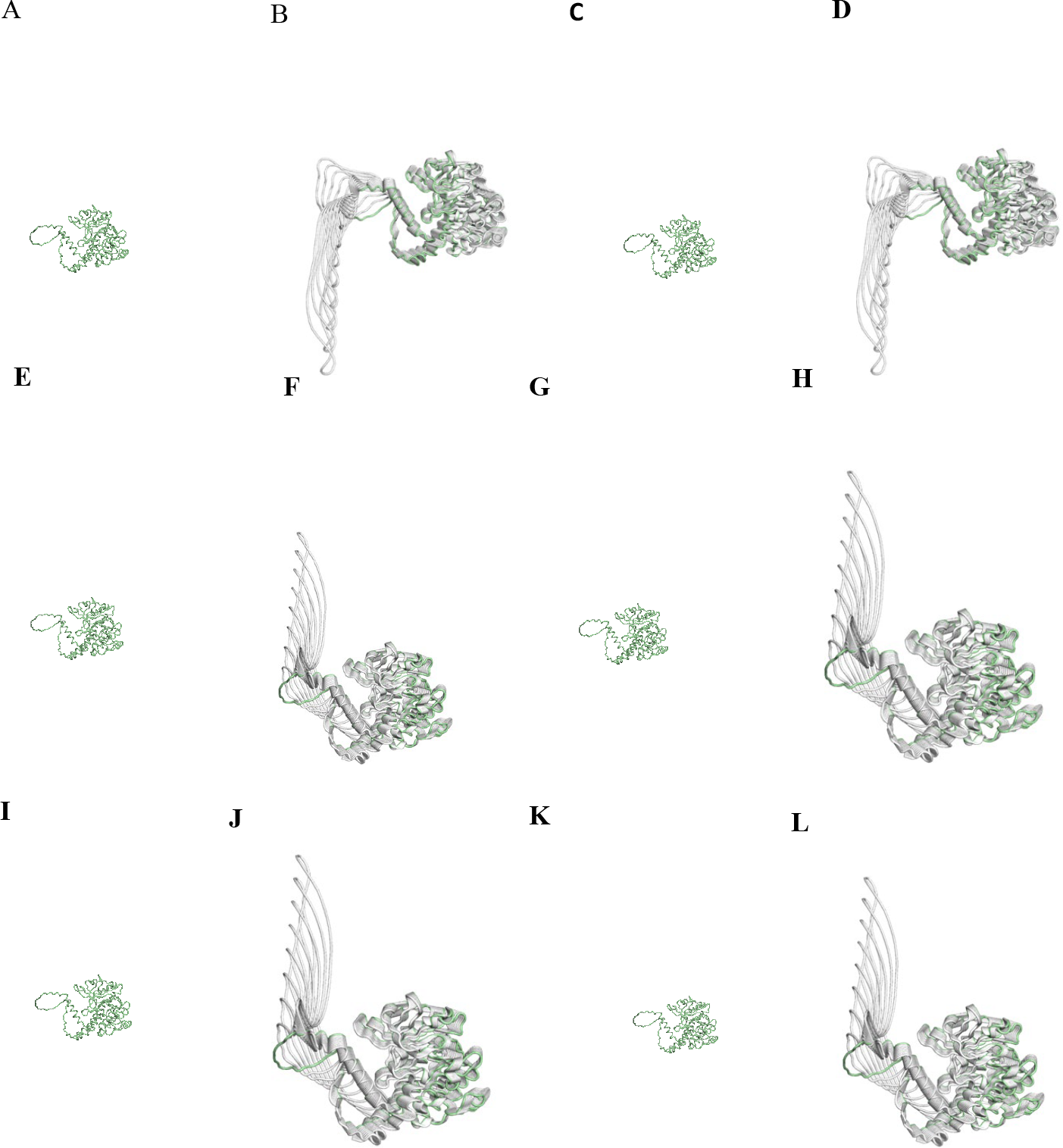
The trajectory representation for the first non-trivial mode of the molecule motion of WT and mutated structures of FLT3 protein based on the Normal Mode Analysis. **(a-f)** and **(a1-f1)** Gif and steady state images of FLT3-WT, FLT3-Y842C, FLT3-Y842F, FLT3-ITD, FLT3-ITD-Y842C and FLT3-ITD-Y842F structures of FLT3 protein.

## 4. Discussion

Proteins are highly dynamic in nature, which is directly related to their conformational stability. Protein conformational stability has a vital role in regulating the protein’s function, which is considered as one of the main features of a protein, and its three-dimensional folded nature can reveal the organism’s functional capabilities and phenotype. Sometimes, protein conformational stability will be altered by mutations, leading to the protein’s structural alteration. The modified protein structures will go through several post-translational modifications, which are the main hallmarks of several diseases, including cancer. In cancers, the altered structure will lead to assorted protein functions by various mechanisms, including activating the protein function in oncogenes or inactivating function in tumor suppressors. All of these will promote cancer formation and a cancer phenotype [34, 58-61]. So, understanding the dynamic behavior of proteins is very important in structural biology for obtaining an indepth understanding of the molecular mechanism of diseases. This knowledge can help us to understand clinically relevant information and discover more effective therapeutics against diseases.

The proposed work highlights the characteristic behaviors of FLT3 protein when it carries a point mutation of C or F in position 842 in the kinase domain, ITD mutation in JM region (REYEYDL amino acid residue duplication from 595-601 (denoted the W51 ITD mutation) and also combination mutation of ITD (FLT3-ITD) with point mutation of C or F in Y842 region in the kinase domain of FLT3 protein ((FLT3-ITD-Y842C and FLT3-ITD-Y842F) by computational study. The point mutation of C at the TKD (Y842C) in FLT3-WT and F mutation at the TKD (Y842) of the FLT3-ITD mutated protein have already been characterized previously [15,21]. But the point mutation to F in FLT3-WT and point mutation to C in FLT3-ITD have not been studied well. Here we endeavor to comprehend the characteristic behaviors of these mutations of FLT3 protein through computational studies including MD simulation, cluster analysis, and machine learning techniques for using the further therapeutic identification against this mutation. Before incorporating the computational techniques, we first modeled the FLT3-ITD protein structure by homology modeling techniques using SWISS-MODEL server. After that, we introduced the point mutation of C or F amino acid residues in position 842 in FLT3-WT and in position 842 in FLT3-ITD structures of the FLT3 protein by Pymol software. Herein, we generated 4-point mutated structures of FLT3, which included FLT3-Y842C and FLT3-Y842F based on FLT3-WT and FLT3-ITD-Y842C and FLT3-ITD-Y842F based on FLT3-ITD structure. To understand the dynamic behavior and conformational stability of the WT and mutated structures of FLT3 protein, we performed computational techniques on six structures, which included four mutated structures of FLT3 (FLT3-Y842C, FLT3-Y842F, FLT3-ITD-Y842C, and FLT3-ITD-Y842F) along with its WT (FLT3-WT) and FLT3-ITD protein structures. Based on all the six structures, first we performed the computational technique was MD simulation analysis. In this study, we mainly focused on the structural deviation (RMSD), fluctuation (RMSF), structural compactness (Rg), intra-H-bond interactions, and solvent-accessible surface area (SASA) of WT and mutated structure of FLT3 protein. In RMSD, RMSF, Rg, and SASA analysis studies, we usually focus on the lowest values, and in the H-Bond interaction analysis study, we examine the highest values relevant to the protein structures. All these values indicate the protein structures’ good optimization, conformation stability, and compactness nature. From the analysis of RMSD (**Figure 3**) values of WT and mutated structures of FLT3 protein, we got the average values were 0.71nm, 0.80nm, 0.79nm, 0.73nm, 0.72nm and 0.75nm for FLT3-WT, FLT3-Y842C, FLT3-Y842F, FLT3-ITD, FLT3-ITD-Y842C and FLT3-ITD-Y842F respectively. When we compare the average RMSD values of FLT3-WT with RMSD values of the mutated structure of the FLT3 protein, we can learn some exciting things like that, when a point mutation occurs in the FLT3-WT with C or F in position 842, the protein will undergo higher structural deviation. However, when the duplication of seven amino acid residues occurs between 595-601 in FLT3-WT, the protein does not exhibit a significant deviationin contrast to the point mutation of FLT3-WT. However, these duplications affect the entire protein’s normal function. A point mutation in Y842 to C in the FLT3-ITD structure results in a greater reduction in deviation compared to FLT3-ITD, whereas a point mutation to F in position 842 of the FLT3-ITD structure, the structure again represents a higher deviation compared to the ITD mutated structure. Based on all the RMSD values of all the structures, we can conclude that point mutated structures of FLT3-Y842C, FLT3-Y842F, FLT3-ITD, FLT3-ITD-Y842C, and FLT3-ITD-Y842F have been deviated when compared to FLT3-WT. These deviations of the structures may change their conformational states and the normal dynamic nature of the protein.

An assessment of the overall fluctuations rates (RMSF in **Figure 4**) of amino acid residues of WT and mutated structures (0.25nm, 0.33nm, 0.18nm, 0.23nm, 0.20nm, and 0.29nm) reveals that the overall fluctuations in the point mutated structures of FLT3-Y842C and combination mutation of FLT3-ITD-Y849F shows higher fluctuations in their amino acid residues. This may enable the mutated structures to acquire more flexibility in their structure and to potentially engage with more surrounding biomolecules in their relevant pathway. About the RMSF values of FLT3-Y842F, FLT3-ITD, and FLT3-ITD-Y842C, it can be stated that the protein structure will acquire a rigid form by these mutations. All these mutations will also lead to the cell’s abnormal functionality and cancer phenotype nature. From the compactness nature of the WT and mutated structure (**Figure 5**), we could identify that the (2.32nm, 2.33nm, 2.24nm, 2.36nm, 2 .26nm, and 2.33n), the compactness nature of FLT3-Y842C, FLT3-ITD and FLT3-ITD-Y842F structures exhibits considerable variation (more flexibility), and FLT3-Y842F and FLT3-ITD-Y842C structures having a much lower compactness nature (more rigid form) than FLT3-WT structure. Based on the behavior of the mutated proteins, we can conclude that higher and lower compactness (flexibility and rigid form) nature will alter the normal protein conformational stability by the mutation. Based on the SASA (Figure **6**) comparison of WT and mutated structures (218.62nm, 216.48nm, 211.69nm, 227.21nm, 223.59nm, and 225.58nm) of WT and mutant FLT3 protein, it can be noted that SASA values for FLT3-Y842C and FLT3-Y842F are significantly lower than those of WT. This suggests that the mutated structure is more rigid than WT in solvent, which affects the normal function of the protein. However, when examining the SASA values of ITD (FLT3-ITD) and combination mutation structures (FLT3-ITD-Y842C and FLT3-ITD-Y842F), it becomes apparent that it possesses greater flexibility in the solvent and is capable of interacting with a greater number of molecules in its vicinity. From the comparison study of both mutated structures, we can say that the SASA nature of the mutated structure may lead to a deregulated protein action in the diseased pathways. Coming to the intra-H-bond (**Figure 7)** interaction of WT with mutated structures (274, 270,276,271,280, and 278), we could identify that mutated structures also show a higher and lower number of intra-bond interactions than FLT3-WT. The lowest intra-H-bond interaction reveals that the mutated structure is not conformationally stable, and in other words, the highest H-bond interaction shows the mutated structure is more rigid than FT3-WT. Here, the lowest H-bond interaction values of FLT3-Y842C and FLT3-ITD indicate that these structures are not stable, and other mutated structures show higher intra-H-bond, meaning that structures preserve more rigidity than FLT3-WT. These results also indicate that the point mutation, duplication, and combination mutation will significantly affect the protein’s entire conformational stability, changing the dynamic nature of the FLT3 protein.

To get a deeper understanding of the conformational flexibility of the mutated FLT3 protein structure, we also performed cluster analysis (**Figure 8**) on all the protein structures using their backbone RMSF values. From the cluster analysis, we got the total number of amino acid residues involved in the stabled conformation area of the WT and mutated structure of the FLT3 proteins, which were 223, 176, 327, 270, 286, and 202, respectively (FLT3-WT, FLT3-Y842C, FLT3-Y842F, FLT3-ITD, FLT3-ITD-Y842C and FLT3-ITD-Y842F). From the results, we can identify that when a point mutation C occurs in position 842 of FLT3-WT, the stability of the amino acid residues decreases in the helix and sheets, and the protein goes to a more unstable state, and the stable number of amino acids residue changed from 223 to 176 in FLT3-WT (more flexibility in FLT3-Y842C structure). But when an F mutation gets in the 842 position, a greater number of amino acid residues became more stable in the protein, and the stability of the amino acid residue changed from 223(FLT3-WT) to 327(FLT3-Y842F), which also indicates the rigidity of the mutated protein. The cluster analysis result of FLT3-ITD reflects that the number of stabled amino acid residues is higher in FLT3-ITD mutated structure when a duplication of seven amino acid residues introduced in located between residues 595 and 601 of FLT3-WT protein and the number of stable amino acid residues changes from 223-270. This also indicates the rigidity of the mutated structure. By examining the combination mutation of FLT3-ITD with C (FLT3-ITD-Y842C) and F (FLT3-ITD-Y842F) amino acid residues, it is possible to infer that the number of stable amino acid residues has increased in FLT3-ITD-Y842C (286) when compared to FLT3-WT and FLT3-ITD, this observation further supports the notion that mutated protein is rigid overall. Intriguingly, however, we can determine that the introduction of a F mutation at position 842 (in the FLT3-ITD structure) reduced the number of stable amino acid residues to 202, a value lower than that of FLT3-WT (223) and FLT3-ITD (270), this indicates the overall instability nature of the mutated structure (more flexibility).

To comprehend the dynamic nature of WT and mutated structures of FLT3, we employed an additional computational tool, DynaMut, which provides information on the dynamic behavior of the protein based on the harmonic movement by NMA analysis. The visual representation results by the NMA analysis (**Figure 9)** reveal that the point mutation C in the 842 position of the FLT3-WT does not alter its movement direction, rather, it changes its frequency (energy) in movements. Nevertheless, it is possible to discern that the harmonic movement of the proteins proceeds in opposite directions in all other mutated structures. This analysis led us to identify that point mutation, duplication mutation, and combination mutation ITD with point mutation of F and C will substantially alter the structural conformation stability and dynamic nature of the FLT3 protein.

## 5. Conclusion

The findings derived from the structure-based computational techniques illuminate the characteristic behavior that underlies the impact of specific mutations on the FLT3 protein.

Notably, the point mutations C or F in the kinase region, the internal tandem duplication (FLT3-ITD) mutation, and the combined mutation of ITD and point mutation of C or F in the kinase region of FLT3 protein structure (FLT3-ITD-Y842C and FLT3-ITD-Y842F) significantly alter the protein’s conformational stability, flexibility, and compactness nature. These alterations interfere the natural dynamic behavior and function of the protein and potentially induce a diseased state and promote drug resistance during treatment. This insightful information will aid in the future discovery of more clinically relevant therapeutics targeting the deregulated pathway of the FLT3 protein.

