## Supplementary-File_S1 for "Understanding the characteristic behavior of the wild-type and mutant protein structure of FLT3 protein by computational methods"

```
Matrix: EBLOSUM62 Gap penalty: 2.0 Extend penalty: 2.0 Score: 5261.0 Sequence 1 length:993 Sequence 2 length:1000 Alignment length: 1000 Identity: 993/1000 (99.30%) Similarity: 993/1000 (99.30%) Gaps: 7/1000 (0.70%)
```

```
1   MPALARDGGQLPLLVVFSAMIFGTITNQDLPVIKCVLINHKNNDSSVGKSSSYPMVSESP 60
    ||||||||||||||||||||||||||||||||||||||||||||||||||||||||||||
1   MPALARDGGQLPLLVVFSAMIFGTITNQDLPVIKCVLINHKNNDSSVGKSSSYPMVSESP 60

61  EDLGCALRPQSSGTVYEAAAVEVDVSASITLQVLVDAPGNISCLWVFKHSSLNCQPHFDL 120
    ||||||||||||||||||||||||||||||||||||||||||||||||||||||||||||
61  EDLGCALRPQSSGTVYEAAAVEVDVSASITLQVLVDAPGNISCLWVFKHSSLNCQPHFDL 120

121 QNRGVVSMVILKMTETQAGEYLLFIQSEATNYTILFTVSIRNTLLYTLRRPYFRKMENQD 180
    ||||||||||||||||||||||||||||||||||||||||||||||||||||||||||||
121 QNRGVVSMVILKMTETQAGEYLLFIQSEATNYTILFTVSIRNTLLYTLRRPYFRKMENQD 180

241 CTRLFTIDLNQTPQTTLPQLFLKVGEPLWIRCKAVHVNHGFGLTWELENKALEEGNYFEM 300
    ||||||||||||||||||||||||||||||||||||||||||||||||||||||||||||
241 CTRLFTIDLNQTPQTTLPQLFLKVGEPLWIRCKAVHVNHGFGLTWELENKALEEGNYFEM 300

301 STYSTNRTMIRILFAFVSSVARNDTGYYTCSSSKHPSQSALVTIVEKGFINATNSSEDYE 360
    ||||||||||||||||||||||||||||||||||||||||||||||||||||||||||||
301 STYSTNRTMIRILFAFVSSVARNDTGYYTCSSSKHPSQSALVTIVEKGFINATNSSEDYE 360

361 IDQYEEFCFSVRFKAYPQIRCTWTFSRKSFPCEQKGLDNGYSISKFCNHKHQPGEYIFHA 420
    ||||||||||||||||||||||||||||||||||||||||||||||||||||||||||||
361 IDQYEEFCFSVRFKAYPQIRCTWTFSRKSFPCEQKGLDNGYSISKFCNHKHQPGEYIFHA 420

421 ENDDAQFTKMFTLNIRRKPQVLAEASASQASCFSDGYPLPSWTWKKCSDKSPNCTEEITE 480
    ||||||||||||||||||||||||||||||||||||||||||||||||||||||||||||
```

```
421 ENDDAQFTKMFTLNIRRKPQVLAEASASQASCFSDGYPLPSWTWKKCSDKSPNCTEEITE480

541  NISFYATIGVCLLFIVVLTLLICHKYKKQFRYESQLQMVQVTGSSDNEYFYVDF------ 594
     ||||||||||||||||||||||||||||||||||||||||||||||||
541 NISFYATIGVCLLFIVVLTLLICHKYKKQFRYESQLQMVQVTGSSDNEYFYVDFREYEYD 594

595 -REYEYDLKWEFPRENLEFGKVLGSGAFGKVMNATAYGISKTGVSIQVAVKMLKEKADSS 653
     ||||||||||||||||||||||||||||||||||||||||||||||||||||||||||
601  LREYEYDLKWEFPRENLEFGKVLGSGAFGKVMNATAYGISKTGVSIQVAVKMLKEKADSS 660

654 EREALMSELKMMTQLGSHENIVNLLGACTLSGPIYLIFEYCCYGDLLNYLRSKREKFHRT 713
    ||||||||||||||||||||||||||||||||||||||||||||||||||||||||||||
661 EREALMSELKMMTQLGSHENIVNLLGACTLSGPIYLIFEYCCYGDLLNYLRSKREKFHRT 720

714 WTEIFKEHNFSFYPTFQSHPNSSMPGSREVQIHPDSDQISGLHGNSFHSEDEIEYENQKR 773
    ||||||||||||||||||||||||||||||||||||||||||||||||||||||||||||
721  WTEIFKEHNFSFYPTFQSHPNSSMPGSREVQIHPDSDQISGLHGNSFHSEDEIEYENQKR 780

774 LEEEEDLNVLTFEDLLCFAYQVAKGMEFLEFKSCVHRDLAARNVLVTHGKVVKICDFGLA 833
    ||||||||||||||||||||||||||||||||||||||||||||||||||||||||||||
781 LEEEEDLNVLTFEDLLCFAYQVAKGMEFLEFKSCVHRDLAARNVLVTHGKVVKICDFGLA 840

834 RDIMSDSNYVVRGNARLPVKWMAPESLFEGIYTIKSDVWSYGILLWEIFSLGVNPYPGIP 893
    ||||||||||||||||||||||||||||||||||||||||||||||||||||||||||||
841 RDIMSDSNYVVRGNARLPVKWMAPESLFEGIYTIKSDVWSYGILLWEIFSLGVNPYPGIP 900

894 VDANFYKLIQNGFKMDQPFYATEEIYIIMQSCWAFDSRKRPSFPNLTSFLGCQLADAEEA 953
    ||||||||||||||||||||||||||||||||||||||||||||||||||||||||||||
901VDANFYKLIQNGFKMDQPFYATEEIYIIMQSCWAFDSRKRPSFPNLTSFLGCQLADAEEA 960
```

```
954 YQNVDGRVSECPHTYQNRRPFSREMDLGLLSPQAQVEDS 993
   ||||||||||||||||||||||||||||||||||||||||
961MYQNVDGRVSECPHTYQNRRPFSREMDLGLLSPQAQVEDS 1000
```
